# High prevalence of lipopolysaccharide mutants and R2-Pyocin susceptible variants in *Pseudomonas aeruginosa* populations sourced from cystic fibrosis lung infections

**DOI:** 10.1101/2023.04.26.538445

**Authors:** Madeline Mei, Preston Pheng, Detriana Kurzeja-Edwards, Stephen P. Diggle

**Affiliations:** Center for Microbial Dynamics and Infection, School of Biological Sciences, Georgia Institute of Technology, Atlanta, Georgia, USA; Department of Pediatrics, Division of Pulmonary, Allergy and Immunology, Cystic Fibrosis, and Sleep, Emory University School of Medicine, Atlanta, Georgia, USA; Emory+Children’s Center for Cystic Fibrosis and Airway Disease Research, Emory University School of Medicine, Atlanta, Georgia, USA

**Keywords:** R-pyocin, bacteriocin, *Pseudomonas aeruginosa*, cystic fibrosis, lipopolysaccharide

## Abstract

Chronic, highly antibiotic-resistant infections in cystic fibrosis (CF) lungs contribute to increasing morbidity and mortality. *Pseudomonas aeruginosa*, a common CF pathogen, exhibits resistance to multiple antibiotics, contributing to antimicrobial resistance (AMR). These bacterial populations display genetic and phenotypic diversity, but it is unclear how this diversity affects susceptibility to bacteriocins. R-pyocins, i.e. bacteriocins produced by *P. aeruginosa*, are phage tail-like antimicrobials. R-pyocins have potential as antimicrobials, however recent research suggests the diversity of *P. aeruginosa* variants within CF lung infections leads to varying susceptibility to R-pyocins. This variation may be linked to changes in lipopolysaccharide (LPS), acting as the R-pyocin receptor. Currently, it is unknown how frequently R-pyocin-susceptible strains are in chronic CF lung infection, particularly when considering the heterogeneity within these strains. In this study, we tested R2-pyocin susceptibility of 139 *P. aeruginosa* variants from 17 sputum samples of seven CF patients and analyzed LPS phenotypes. We found that 83% of sputum samples did not have R2-pyocin-resistant variants, while nearly all samples contained susceptible variants. there was no correlation between LPS phenotype and R2-pyocin susceptibility, though we estimate that about 76% of sputum-derived variants lack an O-specific antigen, 40% lack a common antigen, and 24% have altered LPS cores. The absence of a correlation between LPS phenotype and R-pyocin susceptibility suggests LPS packing density may play a significant role in R-pyocin susceptibility among CF variants. Our research supports the potential of R-pyocins as therapeutic agents, as many infectious CF variants are susceptible to R2-pyocins, even within diverse bacterial populations.

**IMPORTANCE:** Cystic fibrosis (CF) patients often experience chronic, debilitating lung infections caused by antibiotic-resistant Pseudomonas aeruginosa, contributing to antimicrobial resistance (AMR). The genetic and phenotypic diversity of P. aeruginosa populations in CF lungs raises questions about their susceptibility to non-traditional antimicrobials, like bacteriocins. In this study, we focused on R-pyocins, a type of bacteriocin with high potency and a narrow killing spectrum. Our findings indicate that a large number of infectious CF variants are susceptible to R2-pyocins, even within diverse bacterial populations, supporting their potential use as therapeutic agents. The absence of a clear correlation between lipopolysaccharide (LPS) phenotypes and R-pyocin susceptibility suggests that LPS packing density may play a significant role in R-pyocin susceptibility among CF variants. Understanding the relationship between LPS phenotypes and R-pyocin susceptibility is crucial for developing effective treatments for these chronic infections.

## INTRODUCTION

Cystic fibrosis (CF) is a genetic disorder primarily characterized by chronic respiratory infections, leading to progressive loss of lung function and increased morbidity and mortality in affected individuals (1). The disease arises from a defective anion pump, known as the cystic fibrosis transmembrane regulator (CFTR), which results in an imbalance of chloride on epithelial cell surfaces (1). This imbalance causes mucus dehydration, making it too thick for proper clearing of airways along the respiratory tract in people with CF. Consequently, this creates an environment that promotes bacterial and fungal infections, which are notoriously challenging to treat.

*Pseudomonas aeruginosa*, a Gram-negative opportunistic bacterium, is the most prevalent bacterial pathogen that colonizes and persistently infects CF lungs (2-7). With a large genome, *P. aeruginosa* can regulate diverse metabolic processes, resist antibiotic treatment, and control various complex social behaviors that aid in its persistence in infections (8-16). Once established in the CF lung, *P. aeruginosa* is extremely difficult to eradicate and often becomes multi-drug resistant (MDR). Although *P. aeruginosa* inherently possesses many antibiotic-resistant mechanisms, strains sourced from chronic lung infections are known to become particularly problematic and resistant to multiple antibiotics over time (17-21). While recent data suggests that the approval and use of the new triple CF therapeutic TRIKAFTA® (elexacaftor/tezacaftor/ivacaftor) has decreased the overall prevalence of *P. aeruginosa* infection, the 2019 CF Foundation Patient Registry Annual Report notes that the prevalence of MDR *P. aeruginosa* infection remains constant (22). In the era of antibiotic resistance, new treatment approaches are crucial against *P. aeruginosa* and other multi-drug resistant pathogens, especially in the case of CF chronic lung infections.

One of the challenges in eradicating *P. aeruginosa* from CF lungs is likely due to the high degree of phenotypic, and genomic diversity that evolves within *P. aeruginosa* populations over the course of infection (6, 18, 19, 23, 24). While this heterogeneity is known to affect antibiotic susceptibility, its impact on alternative therapeutic options, such as bacteriocins and bacteriophage, remains poorly understood (6, 18, 19). As pathogens like *P. aeruginosa* become increasingly difficult to eliminate, antimicrobials such as bacteriocins, may serve as useful alternative therapies in combination with antibiotics and phage for more effective treatment.

Bacteriocins are narrow-spectrum, proteinaceous antimicrobials produced by one strain of bacteria and active against other strains of the same or closely related species (25, 26). *P. aeruginosa* produces various bacteriocins called pyocins, which are grouped based on similarities in their physical and chemical properties, export mechanisms, and mode of action (27-31). Pyocins produced by *P. aeruginosa* are classified as S-, R-, and F-pyocins, each with varying antimicrobial spectra (30-34). R-pyocins are narrow-spectrum, contractile, phage tail-like bacteriocins, further categorized into subtypes R1-R5 based on specificity to different receptors on target cells, however R-types 2-4 are often considered as a single, functional subtype (30, 31, 35-38). The proposed mode of action involves the R-pyocin tail fiber foot binding to a monosaccharide receptor on the lipopolysaccharide (LPS) (specific to each R-subtype), causing the sheath to contract and push the tail spike and core to puncture the outer membrane, resulting in membrane depolarization and eventual cell death (**Fig. S1**) (37, 39-44). R-pyocins have gained interest for their potential to combat *P. aeruginosa* due to their powerful ability to kill target cells, their inability to replicate without a DNA-containing head, and their well-documented mechanism of action (30, 31, 41, 43-45). In this study, we focus on R-pyocins because of their promising features as antimicrobial agents and the need for a better understanding of their effectiveness against clinical strains of *P. aeruginosa*.

The LPS of *P. aeruginosa* not only acts as the receptor for R-pyocins, but importantly plays a significant role in virulence and immune evasion. LPS is composed of three major regions: the lipid A, the core oligosaccharide, and the O-specific antigen (OSA) (46). The core structure is relatively conserved across *P. aeruginosa* strains and acts as a platform for the attachment of various OSA side chains; Both OSA and common polysaccharide antigen (CPA) can be found in *P. aeruginosa* LPS, but their presence and heterogeneity vary among different isolates or variants even within the same strain ((46-51). The core structure of *P. aeruginosa* LPS is essential for maintaining outer membrane integrity and plays a crucial role in bacterial survival, thus it is costly to bacterial fitness to alter. LPS is composed of a conserved lipid A region, anchoring LPS to the outer membrane, and a core oligosaccharide region that extends from lipid A (47, 52-54). This core region serves as the attachment site for the OSA and CPA side chains, (as well as R-pyocins) contributing to the overall heterogeneity of *P. aeruginosa* LPS (38, 47, 54-58). OSA structures can differ in their length, sugar composition, and linkage patterns, which contributes to antigenic variation and immune evasion strategies employed by the bacterium during infection (59-61). The presence and composition of OSA and CPA side chains in *P. aeruginosa* LPS can vary significantly between strains, thus the diverse repertoire of OSA structures can influence the bacterium’s interactions with not only other pathogens in infection, but also the host immune system, allowing for evasion of immune surveillance and facilitating chronic infection establishment in CF patients (59, 62-65). Moreover, certain *P. aeruginosa* isolates have been reported to downregulate or even lose OSA expression during chronic CF infections, contributing to increased adaptability and persistence in the host environment (51, 66-68)

During chronic CF infections, P. aeruginosa faces a challenging host environment characterized by immune responses and antibiotic exposure. In this context, certain variants of *P. aeruginosa* have been observed to downregulate or lose OSA expression (46, 51, 69). This phenomenon is thought to provide a survival advantage by reducing recognition and clearance by the host immune system. By modulating OSA expression, these isolates can persist and adapt more effectively within the CF lung environment, leading to prolonged and difficult-to-treat infections. The downregulation or loss of OSA expression by certain isolates during chronic CF infections further underscores the bacterium’s adaptability and persistence in the host environment (46, 69). Understanding the structural and functional characteristics of *P. aeruginosa* LPS is crucial for developing targeted therapeutic strategies to combat these challenging infections effectively, and particularly when considering LPS-binding antimicrobials such as bacteriophage and R-type bacteriocins. While there have been studies serotyping and R-pyocin typing panels of strains (38, 70, 71), there has not yet been a study examining LPS diversity among variants of the same strain or comparing variants within multiple different strains and the resulting impacts on R-pyocin susceptibility.

In previous research, we tested 20 individual *P. aeruginosa* variants from sputum samples collected from three different individuals with cystic fibrosis (CF) (60 variants total, from 3 *P. aeruginosa* populations). We found a variation in susceptibility to R2-pyocins within these populations (51). Additionally, we isolated and compared the lipopolysaccharide (LPS) phenotypes of three of these variants from the same sputum sample and patient, discovering that each variant had different LPS glycoforms, which likely contributed to their varying susceptibility (51). If R-pyocins are to be considered as a therapy, it is essential to further investigate their efficacy against diverse clinical strains of *P. aeruginosa*, as the prevalence of R-pyocin-susceptible strains in CF chronic lung infections is currently not well understood.

We previously found that R1-pyocin producers make up the majority of R-pyocin-typeable CF *P. aeruginosa* strains, and other studies have shown that R1-type producers are often susceptible to R2-pyocins (38, 51, 70). As a result, we expected a high frequency of *P. aeruginosa* strains isolated from chronic lung infections with LPS mutations conferring R2-pyocin susceptibility. Although we discovered no correlation between LPS phenotype and R2-pyocin susceptibility, most *P. aeruginosa* strains isolated from CF lung infections were sensitive to R2-pyocins. Our findings support the potential use of R-pyocins as alternative anti-pseudomonal treatments in chronic infections.

## RESULTS

### The majority of *P. aeruginosa* populations in CF patients have variants susceptible to R2-pyocins

To determine the prevalence of R2-pyocin susceptible variants among *P. aeruginosa* populations from CF sputum samples, we analyzed up to 10 previously described variants (51) from longitudinal samples of 7 individuals with CF (139 total variants collected from 17 sputum samples or “populations”) using the soft-agar overlay method (**Fig. S2**). For consistency, we used PAO1Δ*rmlC*, an LPS core mutant (38), as an R2-pyocin susceptible control and PAO1 wt as a resistant control in each experiment. Using R2-pyocins from PAO1 without the pf4 prophage (PAO1ΔPf4) (72), we assessed the relative susceptibility of each CF variant and found that (i) each population showed heterogeneity within the population, consistent with previous work using a microtiter plate method (51), and (ii) nearly every *P. aeruginosa* population included variants with some degree of susceptibility to R2-pyocins. In our sample, we found that only 3 out of 7 individuals had sputum containing R2-pyocin resistant variants (**Fig. 1**). We also discovered that most populations contained R2-susceptible variants even across longitudinal sampling from the same patient, although the degree of relative susceptibility and sensitivity to R2-pyocins fluctuated over time (**Fig. 2**). Longitudinal sputum collections in our biobank showed no trend of variants becoming R2-pyocin resistant over a period of approximately 2 years. To further confirm that these results were R2-pyocin dependent, we included lysates from a PAO1ΔPf4ΔR mutant (lacking both the prophage and a functional R2-pyocin) in each assay, ensuring no killing activity when R2-pyocins were absent.

**Figure 1.**
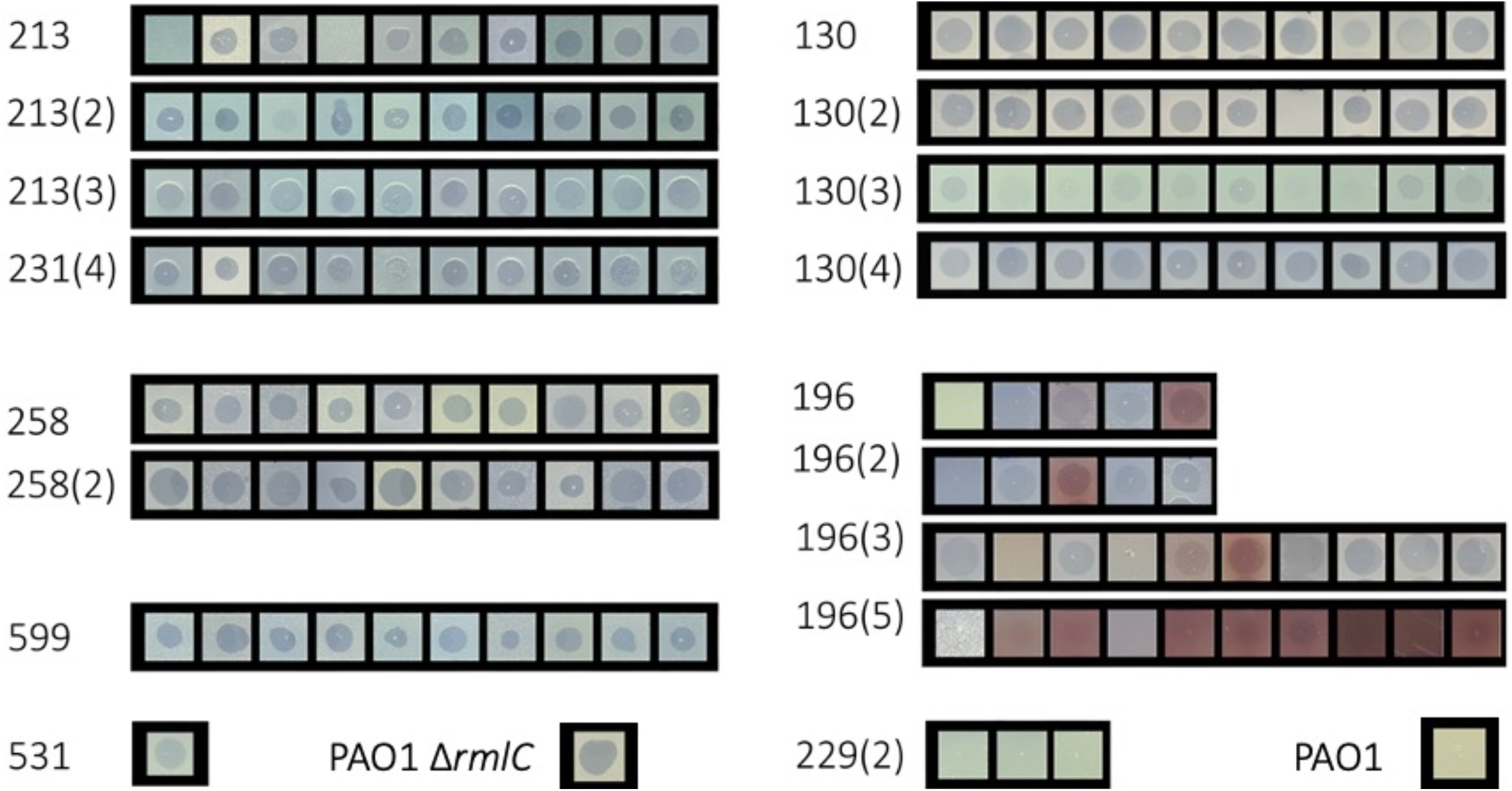
R2-pyocin inhibition zones for multiple CF variants from different patient sputum samples. The inhibition zones caused by R2-pyocin applied to soft agar overlays containing clinical variants from the CF sputum samples listed on the left. PAO1Δ*rmlC* serves as an R2-susceptible control, and PAO1 wt as an R2-resistant control. Red boxes indicate patients with populations containing resistant variants. Up to ten variants per sputum sample were tested. An inhibition zone indicates R2-pyocin susceptibility, while the absence of a visible zone indicates resistance.

**Figure 2.**
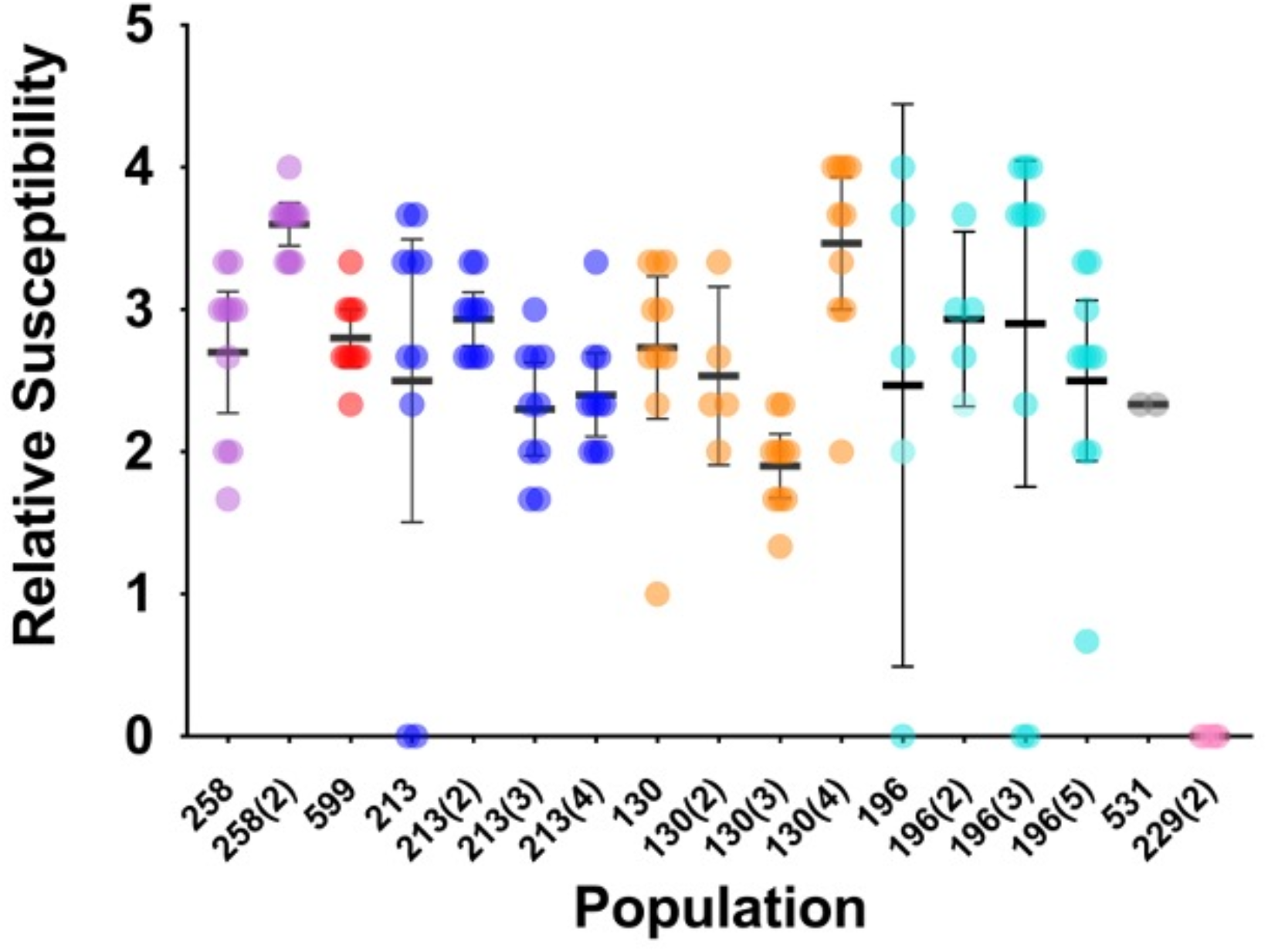
R2-pyocin susceptibility varies among clinical *P. aeruginosa* populations and changes over time. Relative susceptibility to R2-pyocins differs among variants in each population and across populations. Susceptibility heterogeneity is observed across all five patients and within the same patient’s longitudinal samples. There is no trend of strains becoming more resistant to R2-pyocins in longitudinally sampled populations, up to 2 years (consecutive numbers with collection number in parentheses). Two sputum samples contained at least one variant resistant to R2-pyocins (Relative Susceptibility = 0). Relative Susceptibility was determined by counting the number of serial dilutions (including neat) where inhibition zones appeared after R2-pyocins were applied to agar overlays. Means are shown with 95% confidence interval error bars.

### There is a high frequency of LPS mutants and diversity among LPS phenotypes in CF strains of *P. aeruginosa*

To characterize the LPS phenotypes of R2-pyocin susceptible and resistant variants in our biobank, we isolated LPS using a hot-aqueous phenol method (73) and compared the LPS migration profiles of our clinical variants to those of isogenic LPS mutants of our Nottingham PAO1 wild type strain. Out of 139 variants across all populations, we observed that 40% lacked common antigen (CPA or A-band) (**Fig. 3A**) and 76% lacked O-specific antigen (OSA or B-band) (**Fig. 3B**). Additionally, 24% presented altered LPS cores (**Fig. 3C**), as determined by comparing to PAO1. We found that 13 out of the 17 sputum collections sampled contained variants with diverse LPS profiles; approximately 76% of our *P. aeruginosa* populations contained a mixture of variants with varying LPS phenotypes.

**Figure 3.**
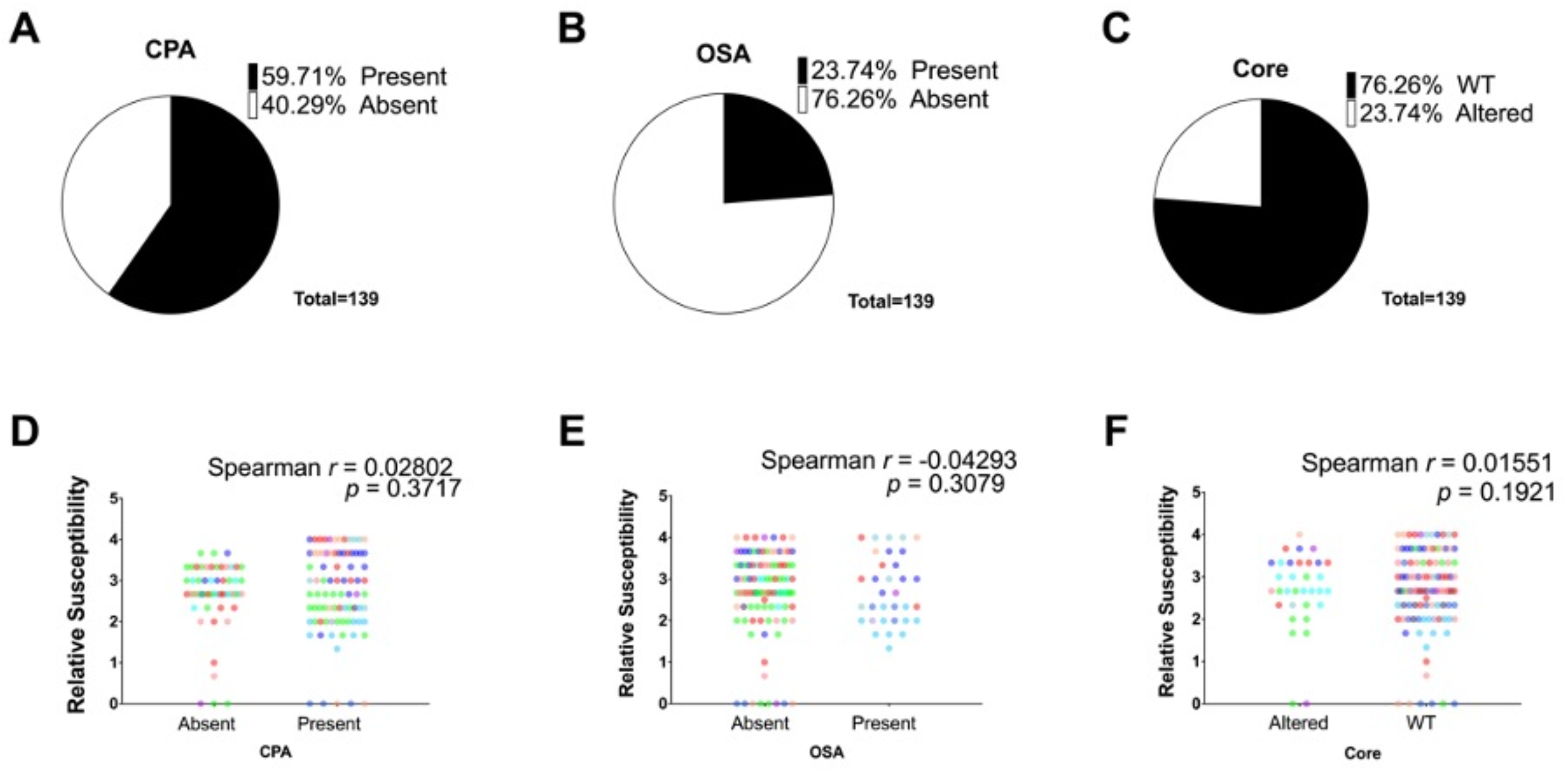
LPS phenotype heterogeneity and lack of correlation with R2-susceptibility. A one-tailed Spearman correlation analysis for each phenotype revealed no correlations between R2-pyocin susceptibility and the absence of (A) common antigen (CPA), (C) O-specific antigen (OSA), or (E) an altered core (p > 0.05). CPA was absent in 42.73% of variants (B), OSA was absent in 76.36%, and (F) 27.27% of variants had altered LPS cores. Each data point represents the average Relative Susceptibility of a variant, and variants are colored based on their longitudinal collection.

To assess the correlation between R2-pyocin susceptibility and specific LPS glycoform phenotypes in CF strains, we conducted a one-tailed Spearman correlation analysis for each phenotype individually. We assumed independence between the three glycoform phenotypes (CPA+ or CPA-, OSA+ or OSA-, WT or altered core) for these analyses, even though some dependencies may exist due to overlapping biosynthetic pathways and the essentiality of the LPS core. We found no significant correlations specifically between the presence or absence of CPA (**Fig. 3D**), OSA (**Fig. 3E**), or an altered core (**Fig. 3F**) (p > 0.05).

We further evaluated the relationship between LPS phenotype and R2-pyocin susceptibility by assessing the LPS as a whole. Using a numbered scale, we assigned an LPS Score of 0 when a variant presented as CPA-, OSA-, and had an altered core; this variant was treated as having 0/3 WT LPS characteristics. A score of 1 was assigned when a variant exhibited one of the three LPS glycoforms as WT (presence of OSA, CPA, or unaltered core), a score of 2 when two of three glycoforms were WT, and a score of 3 when a variant presented CPA+, OSA+, with a WT core. Using this classification system, we also found no correlation with R2-pyocin susceptibility (Fig. 4). However, we did find that three R2-pyocin resistant variants were all CPA- and OSA-. Only 20% of variants from all sputum samples exhibited CPA+, OSA+, and WT LPS cores.

**Figure 4.**
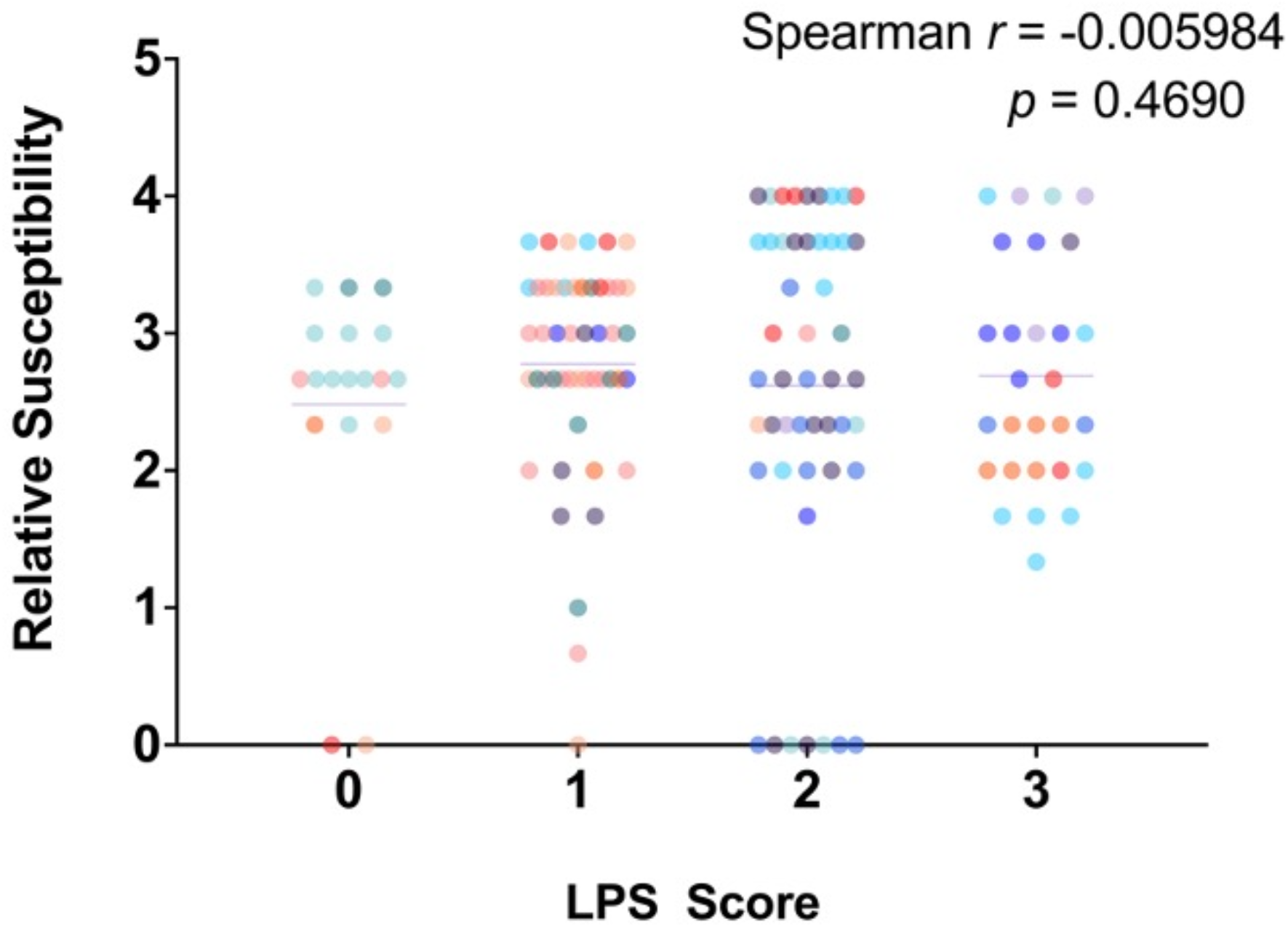
No correlation between combined LPS deficiency phenotypes and R2-pyocin susceptibility. An LPS Score of 0 to 3 was assigned to each variant from each sputum collection to assess the correlation between overall LPS phenotypes and R2-pyocin susceptibility. Variants from each population are shown as data points of the same color. No correlation was found between LPS Score and R2-pyocin susceptibility (p > 0.05).

## DISCUSSION

Cystic fibrosis (CF) chronic lung infections present a significant challenge due to their high antibiotic resistance, difficulty in eradication, and potentially lethal nature (1). *P. aeruginosa*, a common pathogen found in CF lungs, contributes to infection-related morbidity and mortality due to its significant antibiotic resistance and ability to diversify over time (2-7). As a result, it is crucial to explore alternative antimicrobial and therapeutic options for treating these chronic infections.

Bacteriocins, particularly R-pyocins, have recently gained attention for their high potency, narrow spectrum, and anti-pseudomonal properties. However, the efficacy of R-pyocins and other phage-like bacteriocins remains uncertain due to the diverse phenotypic behavior of *P. aeruginosa* in chronic CF lung infections. Previous work demonstrated that *P. aeruginosa* populations display varying susceptibility to R-pyocins, emphasizing the need to better understand the implications of this heterogeneity on treatment outcomes (51). This heterogeneity is particularly important when considering resistant variants in infection, as the prevalence of resistant variants and the benefits of resistance-related phenotypes to survival in chronic infection remain unclear.

In a previous study, we tested the R2-pyocin susceptibility of 60 isolates from 3 patients expectorated sputum samples, but the limited sample size did not provide a comprehensive understanding of R-pyocin sensitivity and resistance behaviors over time or across more CF patients. In the current study, a larger number of variants were tested, collected from multiple longitudinal samples and more individuals with chronic CF lung infections. A total of 139 *P. aeruginosa* variants were collected from 17 expectorated sputum samples (including longitudinal collections) of 7 individuals with CF and were tested for R2-pyocin susceptibility and LPS phenotypes.

The analysis of *P. aeruginosa* populations from CF lung infections revealed that (i) approximately 76% of sputum samples contained *P. aeruginosa* populations exhibiting heterogeneity among LPS phenotypes; (ii) no correlation was found between R2-pyocin susceptibility and the presence or absence of a specific LPS glycoform; and (iii) it was estimated that about 76% of the variants sampled from sputum lacked O-specific antigen, 40% lacked common antigen, and 23% exhibited altered LPS cores.

In previous work, R-pyocin susceptibility testing has typically focused on one variant per sputum collection (38, 70). However, the current study expanded this approach by examining multiple variants from a wide range of sputum samples from various individuals, including some longitudinal samples. This comprehensive analysis revealed that nearly all bacterial populations studied contained susceptible variants (16 out of 17 populations), with heterogeneity in susceptibility observed among the susceptible variants (51). Only about 5% of the tested variants were resistant to R2-pyocins, and these variants were obtained from only 4 sputum samples from 3 patients. No trends in R2-pyocin susceptibility were identified across longitudinal samples, and susceptibility did not cluster by patient, but by longitudinal collection.

The high frequency of R2-pyocin susceptible strains in CF lung infections suggests that (i) there is no strong selective pressure for *P. aeruginosa* to alter LPS in a way that leads to R2-pyocin resistance, and (ii) R2-pyocins could be effective against *P. aeruginosa* for the majority of individuals with CF lung infections, as 94% of individuals in the cohort were infected with R2-pyocin susceptible variants.

Previous studies have shown there is some correlation between OSA serotype and R-pyocin genotype, and though susceptibility to R1, R2, and R5 pyocins were tested for representative strains of varying serotypes, no statistic was provided linking O-antigen serotype with R-pyocin susceptibility (38, 70). Several specific residues on the outer core have been identified as binding factors for each of the 3 functional subtypes of R-pyocins (38) and this implies that O-serotype alone is not the sole determinant of R-pyocin susceptibility as the residues recognized are present among all naturally occurring glycoforms of LPS regardless of serotype. It has not been shown that any of the serotypes have differently structured outer cores, thus in this work, we sought to first look at the presence or absence of the LPS “decorations” (OSA and CPA) and the impacts of variations in these phenotypes rather than correlate serotypes. This study is novel in examining LPS glycoform diversity (i.e. presence or absence of OSA and CPA) among variants of the same strain or comparing variants within multiple different strains and the resulting impacts on R-pyocin susceptibility.

In this study, LPS phenotypes were analyzed based on the presence or absence of the CPA, the OSA, and the altered or PAO1-like (WT) LPS core. No correlations were found between any of the phenotypes or between various combinations of LPS phenotypes and R2-pyocin susceptibility. However, some sputum samples contained variants with similar LPS phenotypes. For example, one sputum collection had all variants exhibiting the phenotype CPA+, OSA+, and presenting a WT core. A high percentage of variants lacked OSA (76%), while more than half maintained the CPA (40%).

The absence of correlation between specific LPS glycoform phenotypes suggests that other LPS-modifying behaviors, such as “packing density”, may be more influential in determining R2-pyocin susceptibility. LPS packing density refers to the arrangement and density of lipopolysaccharide (LPS) molecules on the outer membrane of Gram-negative bacteria, which is also essential for the stability and integrity of the bacterial outer membrane (38, 74-76). Packing density determines how closely the LPS molecules are arranged on the surface of the membrane, which can influence several aspects of bacterial physiology, including the permeability of the outer membrane to antimicrobial agents, the accessibility of OSA side chains, affecting the recognition of bacterial pathogens by immune cells and the production of antibodies, biofilm formation, and the exposure and accessibility of residues targeted by bacteriophage and bacteriocins (including R-pyocins)}(74-76). Other groups have proposed the importance of LPS packing density and the impact on R-pyocin susceptibility (38, 50). One group found several genes responsible for outer membrane lipid homeostasis and LPS packing via TnSeq, including the *mla* and *lpt* transport pathways (50). Though there is some identification of pathways involved in maintaining outer membrane integrity, lipid homeostasis, and LPS arrangement, the regulation of these behaviors remain poorly understood.

In summary, the majority of *P. aeruginosa* strains isolated from CF lung infections appear to be susceptible to R2-pyocins, supporting the potential for R-pyocins as effective antimicrobials. This study also revealed no correlation between LPS phenotype and R2-susceptibility in strains evolved in chronic infection, indicating that further research is necessary to evaluate other LPS-modifying behaviors and packing density in the CF lung environment, as well as the implications of these behaviors for the use of LPS-binding phage and bacteriocins.

## MATERIALS AND METHODS

### Bacterial strains, media, and culture conditions

We collected expectorated sputum samples for this study in previous work from adult CF patients through Emory–Children’s Center for Cystic Fibrosis and Airways Disease Research by the Cystic Fibrosis Biospecimen Laboratory with IRB approval (Georgia Tech approval H18220) (51). We isolated *P. aeruginosa* populations from each sputum sample using selective media (Pseudomonas Isolation Agar, Sigma-Aldrich) before isolating single colonies for further characterization. In total we studied 139 variants sourced from CF chronic lung infections for R2-pyocin susceptibility and LPS phenotype. All *P. aeruginosa* lab strains used in this work are listed and described in **Table 1**. All *P. aeruginosa* strains isolate from CF sputum are listed and described in **Supplementary Dataset 1**. We grew all bacterial cultures in lysogeny broth (LB) medium at 37°C with 200 rpm shaking agitation and used standard genetic techniques for construction of *P. aeruginosa* mutants. The construction of plasmids and generation of mutants are described below and all plasmids and primers used for mutant generation can be found in **Table 1**.

### Generation of mutants

We constructed the PAO1*ΔrmlC* mutant as follows: 600 bp DNA sequences flanking the open reading frames of *rmlC* (PA5164) were PCR amplified using Q5 DNA polymerase (New England Biolabs, Ipswich, MA) (38). We then cloned these sequences into the SphI-XbaI digested suicide vector pDM4 (50) by Gibson assembly using NEBuilder HiFi Assembly master mix (New England Biolabs), transformed into *E. coli* S17-λpir by electroporation and selected on LB agar plates supplemented with 30 μg/mL chloramphenicol. We verified cloned inserts by colony PCR and Sanger sequencing (Eton Bioscience, NC). We introduced the deletion construct into PAO1 wt by electroporation and strains carrying single crossover insertions of the deletion constructs were selected on LB agar plates supplemented with 300 μg/mL chloramphenicol (77, 78). Resistant colonies were cultured in LB without antibiotic and plated on LB agar plates with 0.25% NaCl and 5% sucrose to select for loss of the construct (79). We screened sucrose-resistant colonies for antibiotic sensitivity to ensure loss of the vector sequence and were assessed for presence of the gene deletion by colony PCR and Sanger sequencing of the PCR product. We constructed the PAO1ΔPf4ΔR mutant by deleting the tail fiber and chaperone genes of the R-pyocin gene cassette (PA0620-PA0621) from the PAO1ΔPf4 strain as described previously (51, 72).

### Expression and collection of R-pyocins

We collected R-pyocins in lysates as previously described (51). LB cultures of PAO1, PAO1ΔR (R-pyocin null mutant) (46), PAO1ΔPf4 (prophage mutant) (72), and PAO1Δ*pf4* ΔR were inoculated at 1:100 from overnight planktonic cultures, and grown to mid-logarithmic growth phase (approximately 4 h) in 10 mL of LB media. We then added ciprofloxacin to each culture for a final concentration of 0.1 μg/mL to induce R-pyocin production (51, 80, 81), and cultures were incubated for a further 3 h. Chloroform was used to lyse remaining cells, before centrifuging lysates at ∼3,300 *x g* for 10 min. The R-pyocin-containing supernatants were separated and stored at 4°C.

### Spot assay for Relative R-pyocin Susceptibility

Spot assays to assess R-pyocin activity were conducted as previously described (51). We used overnight cultures of clinical *P. aeruginosa* variants to inoculate 4 mL of cooled soft top (0.4%) agar at an optical density at 600 nm (OD_600_) of 0.01. This mixture was poured onto LB agar plates for an overlay of the indicator strain. We extracted R2-pyocin lysates from PAO1, PAO1ΔPf4 and respective R-pyocin null mutants were vortexed, then serial diluted 10-fold in phosphate buffered saline (PBS) before spotting 5 μL of each dilution onto the soft-top overlay indicator strains. Spots were allowed to dry before incubating plates at 37°C overnight. Clear zones of growth inhibition from PAO1ΔPf4 lysates were used to determine R-pyocin dependent activity against the indicator strains, and we determined relative susceptibility by counting the number of these serial dilutions of lysate indicating inhibition. PAO1 wild type was used as an R2-resistant control strain and PAO1Δ*rmlC* was used as an R2-susceptible control strain to standardize and quality control each experiment (38). We conducted spot assays for R-pyocin susceptibility in triplicate.

### LPS isolation and visualization

We isolated LPS of bacterial cultures and visualized as described by previous methods (73). We used overnight bacterial cultures of 5 mL LB grown at 37°C at 200 rpm shaking. We then diluted overnight cultures 1:10 with LB to measure OD_600_ before making a 1 mL suspension of bacteria adjusted to an OD_600_ of 0.5 and pelleting by centrifugation at 10,600 *x g* for 10 min. The cell pellet was resuspended in 200 μL of 1x SDS-buffer. Suspended bacterial cells were boiled for 15 min before we prepared LPS by hot aqueous-phenol extraction as previously described but without the addition of DNase I and RNase solutions (as deemed optional in the protocol) (73). We visualized samples using 15 μL of LPS preparation on a 4%-20% Tris-glycine mini gel using the Pro-Q Emerald 300 Lipopolysaccharide Gel Stain Kit (ThermoFisher) (73).

### Statistical analysis

We performed all data analysis using Prism (Graphpad software, version 9).

## Data Availability

We have made raw data and code available in the Dryad Digital Repository: https://datadryad.org/stash/share/k9quE9rIzJ-gRXo42tF559U2MEpmXH6wC3emdpinkkM (82).

## Supplemental Material

### Supplementary Dataset 1

Detailed description of bacterial strains, average susceptibility and LPS scoring used in this study.

## Acknowledgements

We thank the Cystic Fibrosis and Airways Disease Research (CF-AIR) at Emory University, for clinical CF populations of *P. aeruginosa*. We also thank Patrick Secor for generously sharing the PAO1ΔPf4 strain with us. This work was supported by the Cystic Fibrosis Foundation by grants (DIGGLE18I0 and DIGGLE20G0),the National Institutes of Health (NIH) for a grant (R01AI153116) and the National Science Foundation (NSF) for a grant (2003721) to SPD. This work was also supported by the NIH Vaccinology Training Program Grant (#T32AI074492) to MM.

## Figure legends

Table 1. Strains, primers and plasmids used in this study.

---

**Figure S1.**
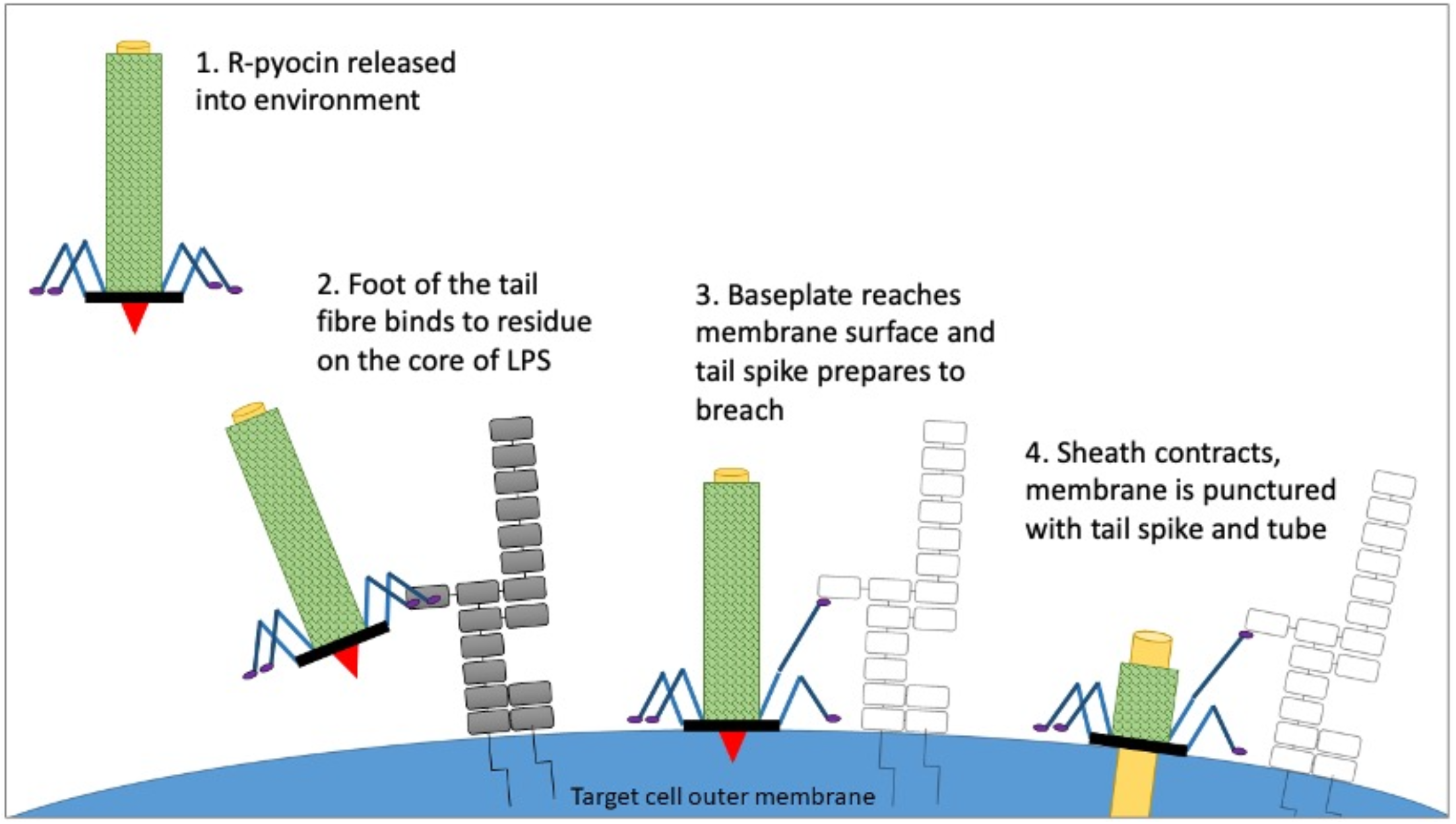
R-pyocin binding to LPS receptor and puncturing target cell membrane. When released into the environment by producing cells, R-pyocins bind to target cells by recognizing core residues of the lipopolysaccharide (LPS) on the outer cell membrane. After tail fiber binding, the baseplate dissociates, initiating sheath contraction and driving the iron-tipped tube through the cell surface, killing the target bacterium.

**Figure S2.**
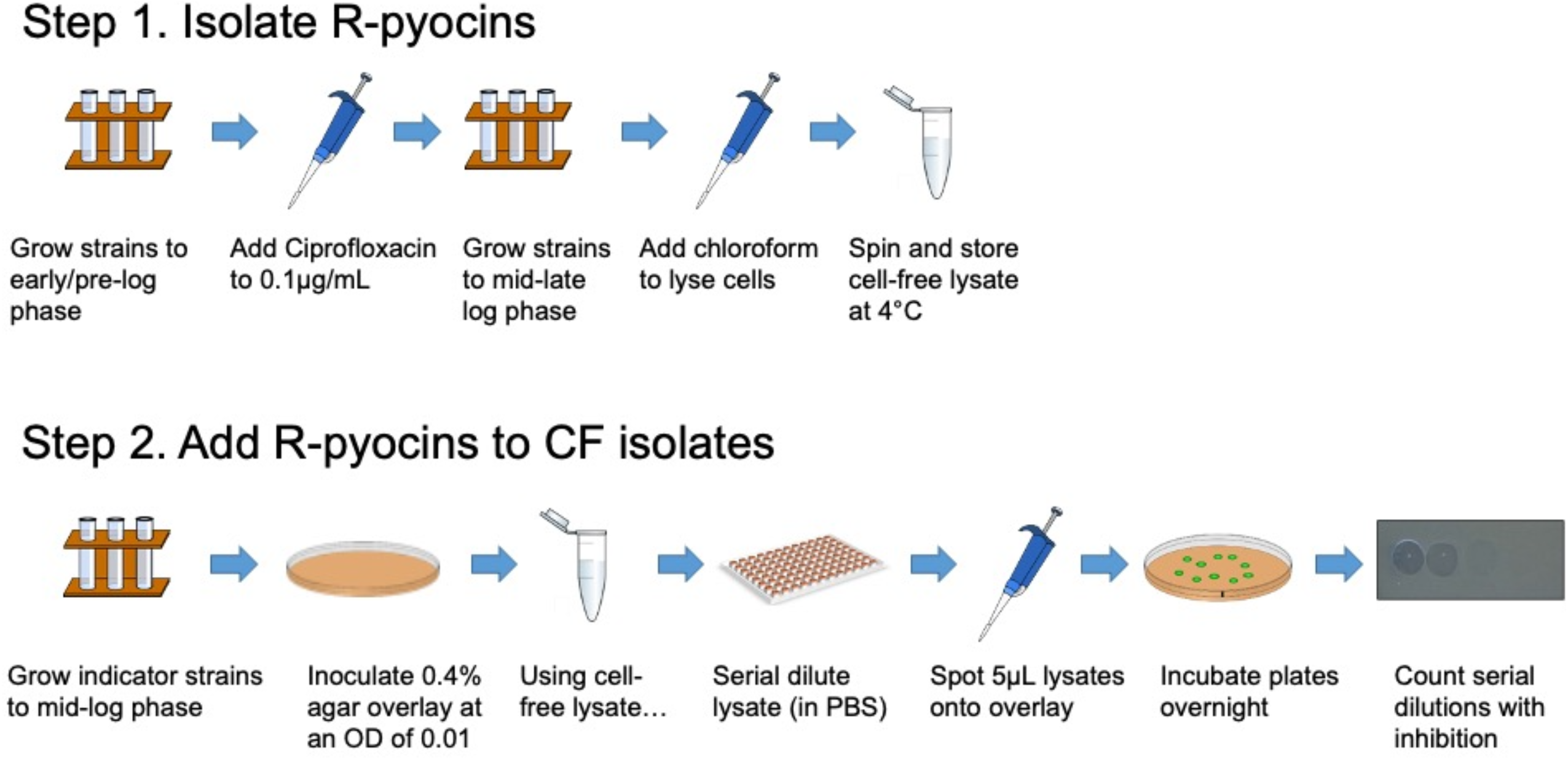
R2-pyocin lysate collection and susceptibility testing procedure. R-pyocins are isolated from *P. aeruginosa* by growing strains to early logarithmic growth, adding sub-minimal inhibitory concentration (MIC) ciprofloxacin, and lysing cells with chloroform after several hours for maximum R-pyocin production. Cultures are then centrifuged, separating into phases, allowing for the collection of cell-free R-pyocin-containing supernatant (lysates) to be stored. R-pyocin lysates are serially diluted and spotted onto soft agar overlays to test the susceptibility of strains of interest.

